# Circulating PD-1^hi^ CXCR5- and CXCR5+ CD4 T cells are elevated in patients with newly diagnosed Giant Cell Arteritis and predict relapse

**DOI:** 10.1101/2024.05.12.593745

**Authors:** Irene Monjo, Beatriz Nieto-Carvalhal, Mariela Uyaguari, Sara García-Carazo, Alejandro Balsa, Eugenio de Miguel, María-Eugenia Miranda-Carús

## Abstract

**BACKGROUND:** Giant cell arteritis (GCA) is a large/medium-vessel granulomatous vasculitis, and the PD-1/PD-L1 coinhibitory pathway seems to be implicated in its pathogenesis. CD4 T cells expressing high PD-1 levels, CD4+CXCR5-PD-1^hi^ peripheral helper (Tph) and CD4+CXCR5+PD-1^hi^ follicular helper T cells (Tfh), are key mediators of autoimmunity. Their frequencies are elevated in the peripheral blood of subjects with several autoimmune conditions but have not been investigated in GCA. Our objective was to study the frequency of circulating Tph (cTph) and Tfh (cTfh) in patients with newly diagnosed GCA (nGCA).

**METHODS:** Prospective, non-interventional study on consecutive patients referred to our ultrasound GCA fast-track clinic over a period of 24 months. Peripheral blood was drawn immediately upon initial diagnosis. For each patient, an age and gender-matched healthy control (HC) was included. PBMCs isolated by Ficoll-Hypaque were examined by cytometry. Patients were subsequently treated with standard therapy according to the updated 2018 EULAR recommendations.

**RESULTS:** 65 nGCA patients were included. As compared with HC, nGCA patients presented at baseline with an increased frequency of cTph and cTfh cells. Among the 46 patients who could be followed up for 12 months, 19 experienced a relapse. The baseline frequency of cTph and cTfh cells had been significantly lower in patients who relapsed as compared with those who did not. A cTph cell frequency <0.56 predicted relapse with a sensitivity of 90% and specificity of 93%.

**CONCLUSION:** nGCA patients demonstrate increased baseline cTph and cTfh cell frequencies. Lower baseline proportions of cTph and cTfh cells associate with relapse.

---

Giant cell arteritis (GCA) is a large- and medium-vessel granulomatous vasculitis involving the aorta and its branches [1]. Despite intensive research its pathogenesis is not well understood, although several clinical and experimental observations point to a participation of the PD-1/PD-L1 coinhibitory pathway [2–6]. The vessel walls show a transmural infiltrate spanning across the intima, media and adventitia layers; it contains activated PD1+CD4+ T cells [2,3] as well as macrophages that tend to fuse into multinucleated giant cells and demonstrate an insufficient PD-L1 expression [2]. Of note, new onset GCA has been described in cancer patients treated with anti-PD-1 monoclonal antibodies [4–6]. Also, immunocompromised mice with engrafted human arteries develop vasculitis after reconstitution with lymphocytes from GCA patients [2]; furthermore, blocking anti-PD-1 antibodies accelerate the disease course and augment the numbers of PD-1+ T cells in the inflamed vessels [2].

Two populations of CD4+ T cells expressing high levels of PD-1 (PD-1^hi^) are key actors in autoimmune conditions: CD4+CXCR5-PD-1^hi^ peripheral helper T cells (Tph) and CD4+CXCR5+PD-1^hi^ follicular helper T cells (Tfh). Tph cells, initially described at the inflamed synovium [7] and peripheral blood of patients with Rheumatoid Arthritis (RA) [7,8], seem to play a pivotal role in the local inflammatory infiltrates of RA and also of other immune-mediated diseases [9] but their possible participation in GCA has not been investigated. On the other hand, Tfh cells dwell in the germinal centers of lymphoid organs [10,11] and are required for the development of autoimmunity [10,11]; furthermore, their circulating counterparts (cTfh) are elevated in patients with several autoimmune conditions [8–15]; of note, previous reports on the frequencies of circulating CD4+CXCR5+ T cells in GCA [16,17] have not described their PD-1 expression. Therefore, our objective was to study the frequencies of circulating CD4+CXCR5-PD- 1^hi^ Tph (cTph) and CD4+CXCR5+PD-1^hi^ Tfh (cTfh) cells in the peripheral blood of patients with newly diagnosed GCA (nGCA), their possible variation along the disease course, and their relation with the clinical outcome.

## PATIENTS AND METHODS

### Ethics Statement

The study was approved by the Hospital La Paz - IdiPAZ Ethics Committee, and all subjects provided written informed consent according to the Declaration of Helsinki.

### Patients

This is a prospective non-interventional study performed on consecutive patients referred to our ultrasound (US) GCA fast track clinic, in whom newly diagnosed GCA (nGCA) was clinically confirmed over a period of 24 months (Table 1).

**Table 1.** Clinical Features of nGCA patients.

|  | **Clinical features (all)** | **Clinical features of patients with 1 year follow-up (n=46)** |  | **Relapse rate at 1yr attending to US/ clinical subgroups** |
| --- | --- | --- | --- | --- |
|  | **nGCA-all n=65** | **nGCA-nr n=27** | **nGCA-r n=19** | **Total Relapses n=19/46 (41%)** |
| Gender: F, n (%) | 37 F (57) | 16 F (59) | 12 F (63) |  |
| Age (yrs): median (IQR) | 82 (74-86) | 82 (74-86) | 79 (72-85) |  |
| Cranial, n (%) | 37 (57) | 14 (52) | 11 (58) | 11/25 (44) |
| Large Vessel, n (%) | 12 (18) | 5 (18) | 4 (21) | 4/9 (44) |
| Mixed, n (%) | 16 (25) | 8 (30) | 4 (21) | 4/12 (30) |
| Polymyalgia Rheumatica, n (%) | 22 (34) | 8 (30) | 9 (47) | 9/17 (53) |

Our fast-track clinic provides rapid clinical and ultrasound evaluation of patients with suspected GCA. The patients were referred from primary care and from La Paz University Hospital medical departments (Rheumatology, Emergency, Internal Medicine, Ophthalmology and Neurology). In most cases Colour Doppler US (CDUS) examination was performed within the first 24 h after referral, with a maximum of 72 h, and patients were naïve for glucocorticoids. Patients with autoimmune diseases other than GCA, cancer or concomitant infections were not included. Peripheral blood was drawn immediately upon initial diagnosis and after obtaining written informed consent. For each patient, an age and gender-matched healthy control (HC) was also studied. Subsequently, patients received standard monotherapy with glucocorticoids following the updated 2018 EULAR recommendations for the management of large vessel vasculitis [18]. In brief, GCA patients were initially treated with 0.7-1 mg/kg/day of prednisone. Tapering of prednisone treatment started after 2-4 weeks, based on normalization of clinical signs and symptoms accompanied by normalization of the erythrocyte sedimentation rate (ESR) and/or C-reactive protein (CRP). Patients were closely monitored following the standardized protocol in our GCA clinic. This includes monthly visits during the first 3 months and quarterly visits during the first 2 years, which allows early detection of relapses defined as per 2018 EULAR recommendations [18]; symptoms plus: elevation of acute phase reactants and/or positive US.

### Colour Doppler Ultrasound (CDUS)

CDUS was performed on the temporal superficial arteries (common temporal, frontal and parietal branches) and extracranial arteries (bilateral common carotids, axillaries and subclavian arteries). The presence or absence of a halo sign [19] and the maximum value of the intima-media thickness (IMT) were recorded in all examined vascular segments. The IMT was measured in the longitudinal view in the wall distal to the US probe and in the area with greatest wall thickness without an atherosclerotic appearance. The cutoff values for IMT, measured from the luminal side of the intima to the adventitia, were: ≥0.34mm for the frontal and parietal branches, ≥0.42mm for the common trunks of the temporal arteries, and ≥1mm for the axillary, subclavian and carotid arteries [20–23]. We defined a CDUS as positive when at least one artery showed a halo sign and its thickness was equal to or greater than the established cut-off point. Patients with isolated involvement of the carotid arteries were excluded to improve the diagnostic validity of the study. An increased IMT in carotid arteries namely lacks the specificity for GCA compared with atherosclerosis as recently shown [23]. Upon diagnosis, patients were classified into US defined phenotypes: cranial (C) large vessel (LV) or mixed (Mx).

Information on specific ultrasound machines and other ultrasound additional information has been previously published [24]. In brief, a high-quality US machine with linear high-frequency probes was employed (Mylab X8 system, Esaote, Genoa, 2017) with the following settings: For temporal arteries, a 12-25 MHz probe was used with a gray scale frequency of 24 MHz, a Doppler frequency of 12.5MHz, an adjusted color gain and a pulse repetition frequency (PRF) of 1.9kHz. For extracranial arteries, a 4-15MHz probe was used, with a gray scale frequency of 15 MHz, a Doppler frequency of 4.5 MHz, an adjusted color gain, and a PRF of 3.0 kHz [25]. The examinations were performed by two expert rheumatologists (E.D.M. and I.M.H.).

### Temporal artery biopsy (TAB)

TAB was not routinely performed. We only request it in doubtful cases, such as patients with a positive CDUS but with low pretest probability or cases where the halo sign is observed in only one arterial branch, attending to the 2018 EULAR recommendations [18].

### Flow Cytometry

The frequency and phenotype of cTph and cTfh cells was assessed by flow cytometry of freshly isolated PBMCs. Events were acquired in a FacsCelesta flow cytometer with FacsDiva software (BD Biosciences) and analyzed using FlowJo 10.8.1 software. Fluorochrome-conjugated mAbs were used to examine the expression of CD3 (BD, clone UCHT1), CD4 (BD RPA-T4), CD45RA (BD L48), CD45RO (BD UCHL1), CXCR5 (BD RF8B2), ICOS (eBioscience ISA-3) and PD-1 (BD EH12.1). The detailed gating strategy for Tfh and Tph cells was applied as previously published by our group [8].

### Statistical Analysis

Comparison between groups was by Mann-Whitney or Kruskal-Wallis test. When appropriate, Bonferroni correction for multiple comparisons was applied. Paired samples were analyzed with the Wilcoxon test. Correlations were analyzed using Spearman’s rank correlation coefficients. Logistic regression was applied to verify the independent associations between immunological and clinical parameters. All analyses were performed using Prism version 10.2.2 software for macOS (GraphPad, California, USA).

## RESULTS

### Baseline Frequencies of cTph and cTfh cells in nGCA patients

The gating strategy for cTph and cTfh cells was applied as we previously described [8]. As compared with HC (n=65), nGCA patients (n=65) demonstrated significantly increased baseline frequencies of circulating CXCR5-PD-1^hi^ (cTph) and ICOS+CXCR5-PD-1^hi^ (ICOS+cTph) CD4+ T cells (Fig.1A-C). In addition, significantly increased frequencies of CXCR5+PD-1^hi^ (cTfh) and ICOS+CXCR5+PD-1^hi^ (ICOS+cTfh) CD4+ T cells were also observed (Fig.1A,B,D). The baseline frequency of CXCR5+ or CXCR5-cells among circulating CD4 + T lymphocytes was not different in nGCA patients (n=65) as compared with HC (n=65) (Fig. S1).

**Figure 1.**
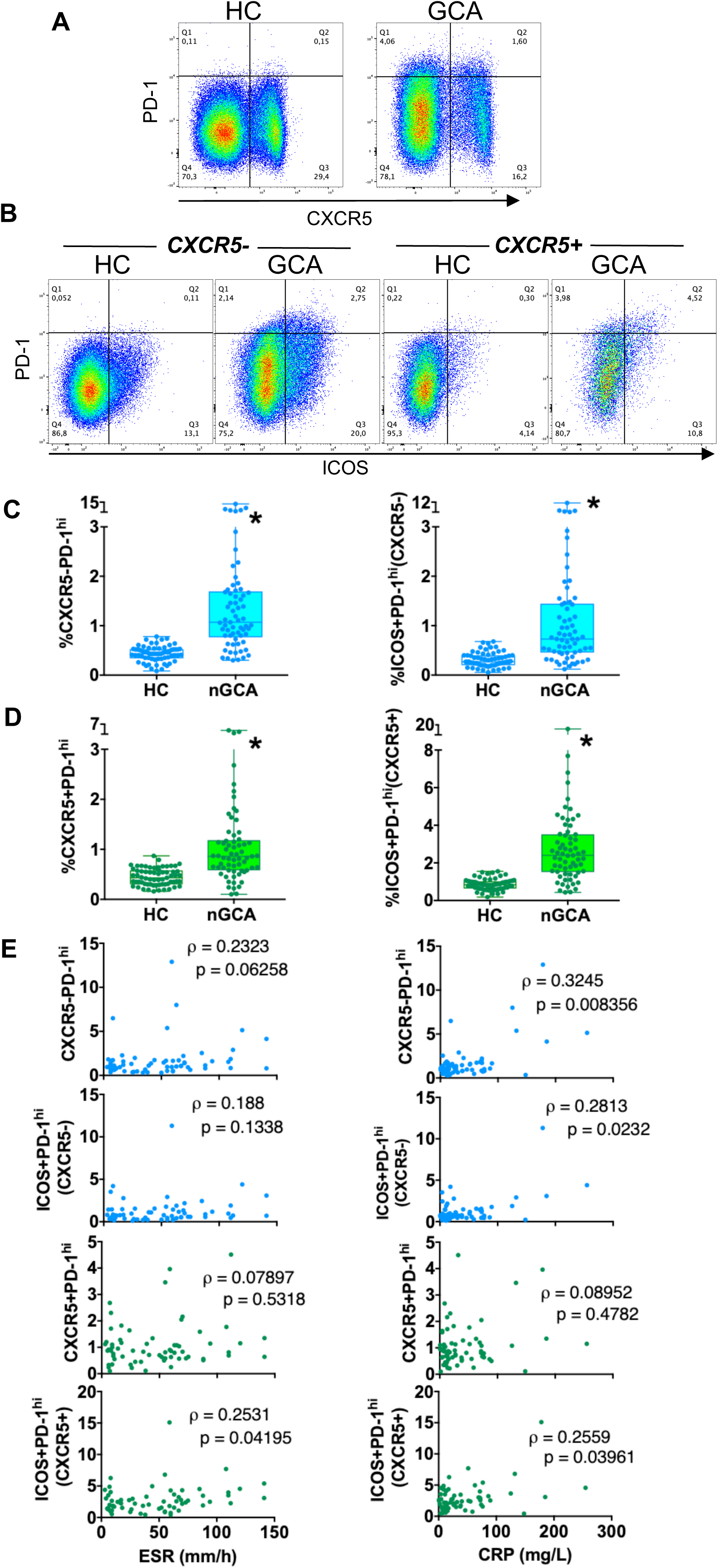
Frequency of cTfh and cTph cells in nGCA patients. **A**. **Representative dot plots** of CXCR5 and PD-1 expression on cells gated for CD3 and CD4 expression. **B. Representative dot plots** of ICOS and PD-1 expression on cells gated for CD3, CD4 and absence (left) or presence (right) of CXCR5 expression. **C**. **Frequency of cTph** cells (CD4+CXCR5-PD-1^hi^) and **ICOS+ cTph** cells (CD4+CXCR5-PD-1^hi^ICOS+). **D**. **Frequency of cTfh** cells (CD4+CXCR5+PD-1^hi^) and **ICOS+ cTfh** cells (CD4+CXCR5-PD-1^hi^ICOS+). **E**. Correlation between ESR or CRP and the frequencies of cTph, ICOS+cTph, cTfh and cTfh (Spearman rho). In **A-D**, box and whiskers plots represent the median, interquartile range, maximum and minimum values; the number of patients per group are expressed in the text and in table 1. For each patient, an age and gender-matched healthy control (HC) is represented. *p<0.05 vs HC, Mann-Whitney test.

The baseline cTph and ICOS+cTph cell frequencies demonstrated a weak correlation with the baseline CRP but not with the baseline ESR. In addition, the baseline ICOS+cTfh cell frequency was weakly correlated with both the baseline CRP and ESR (Fig 1E).

### Baseline frequencies of cTph and cTfh cells in nGCA patients attending to ultrasound or clinically defined phenotypes

The baseline frequencies of cTph, ICOS+cTph, cTfh or ICOS+cTfh cells were not significantly different among patients with different ultrasound defined phenotypes [cranial (n = 37), large vessel (n= 12) or mixed (i.e., cranial plus large vessel) (n = 16)] (Fig.2, A-D), or between patients with (n= 22) or without (n= 43) associated polymyalgia rheumatica (PMR) (Fig.2, A-D).

**Figure 2.**
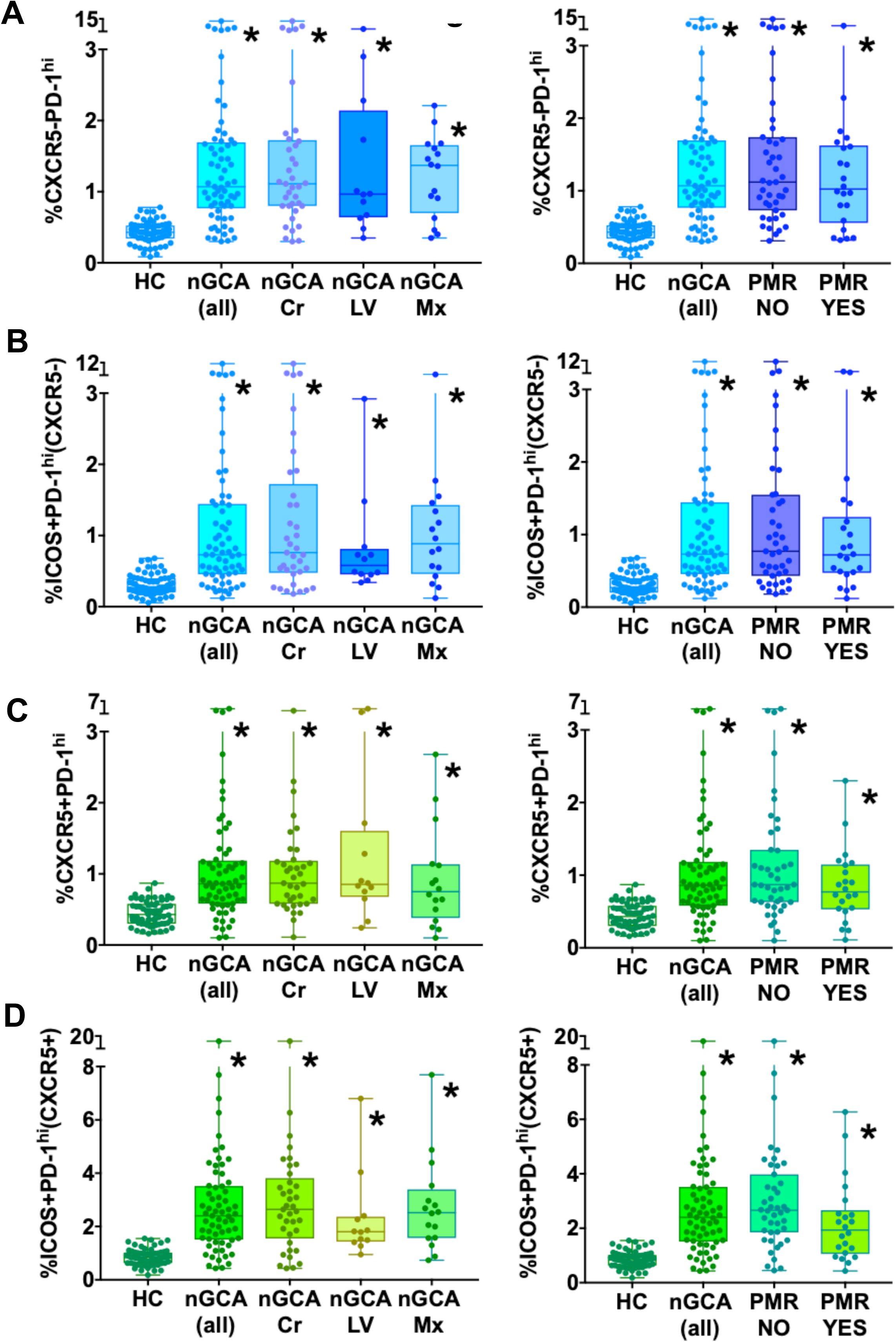
Frequency of cTph and cTfh cells in nGCA patients attending to the ultrasound defined subgroups [cranial (C), extracranial (EC), mixed (Mx)] and clinically defined subgroups [associated or not with polymyalgia rheumatica (PMR)]. **A**. Frequency of cTph; **B**. Frequency of ICOS+cTph; **C**. Frequency of cTfh; **D**. Frequency of ICOS+cTfh. Bars represent the median and interquartile range; whiskers represent the maximum and minimum values. *p<0.05 vs HC, Kruskal-Wallis test with Bonferroni correction.

### Relation of the baseline cTfh or cTph frequencies with the clinical outcome

We next sought to examine the possible relation of the baseline cTph and cTfh cell frequencies with the occurrence of relapse in the first 12 months after diagnosis. A total of 46 nGCA patients completed a 12 month follow-up; 19 of them (41.3%) had experienced a relapse during that time period (Table 1). These were 18 minor relapses and one major relapse as defined by EULAR recommendations [18]. The median time from baseline to relapse was 5 months (IQR 2-10) and the median daily prednisone dose at relapse was 15 mg (IQR 7.5-20). Patients who relapsed in the first 12 months had demonstrated significantly lower baseline frequencies of cTph (Fig.3A) [Odds Ratio (OR) 51 (95% confidence interval, 22-110)], ICOS+cTph (Fig.3B) [OR 15 (6-40)], cTfh (Fig.3C) [OR 4 (1.8-10)] and ICOS+cTfh (Fig.3D) [OR 8 (3-14)] as compared with the group of nGCA patients who had not relapsed, and multiple logistic regression indicated that this was independent of baseline ESR or CRP values, age, gender, US defined subgroup or presence of PMR. The Receiver Operating Characteristic (ROC) analysis demonstrated an area under the curve (AUC) of 0.96 (95% CI 0.91-1.0), p<0.0001, for the association of cTph with relapse; 0.93 (0.87-1.0), p<0.0001 for ICOS+cTph; 0.85 (0.74-0.96) for cTfh and 0.89 (0.80-0.98) for ICOS+cTfh (Fig. 1A-D). The optimal cutoff value for predicting relapse at 12 months was a cTph cell frequency below 0.56 (sensitivity 90%, specificity 93%). Additional cutoff values for relapse were ICOS+cTph <0.37 (S 73.3%, Sp 86.4%), cTfh <0.56 (S 68%, Sp 85%) and ICOS+cTfh<0.61 (S 74%, Sp 93%).

**Figure 3.**
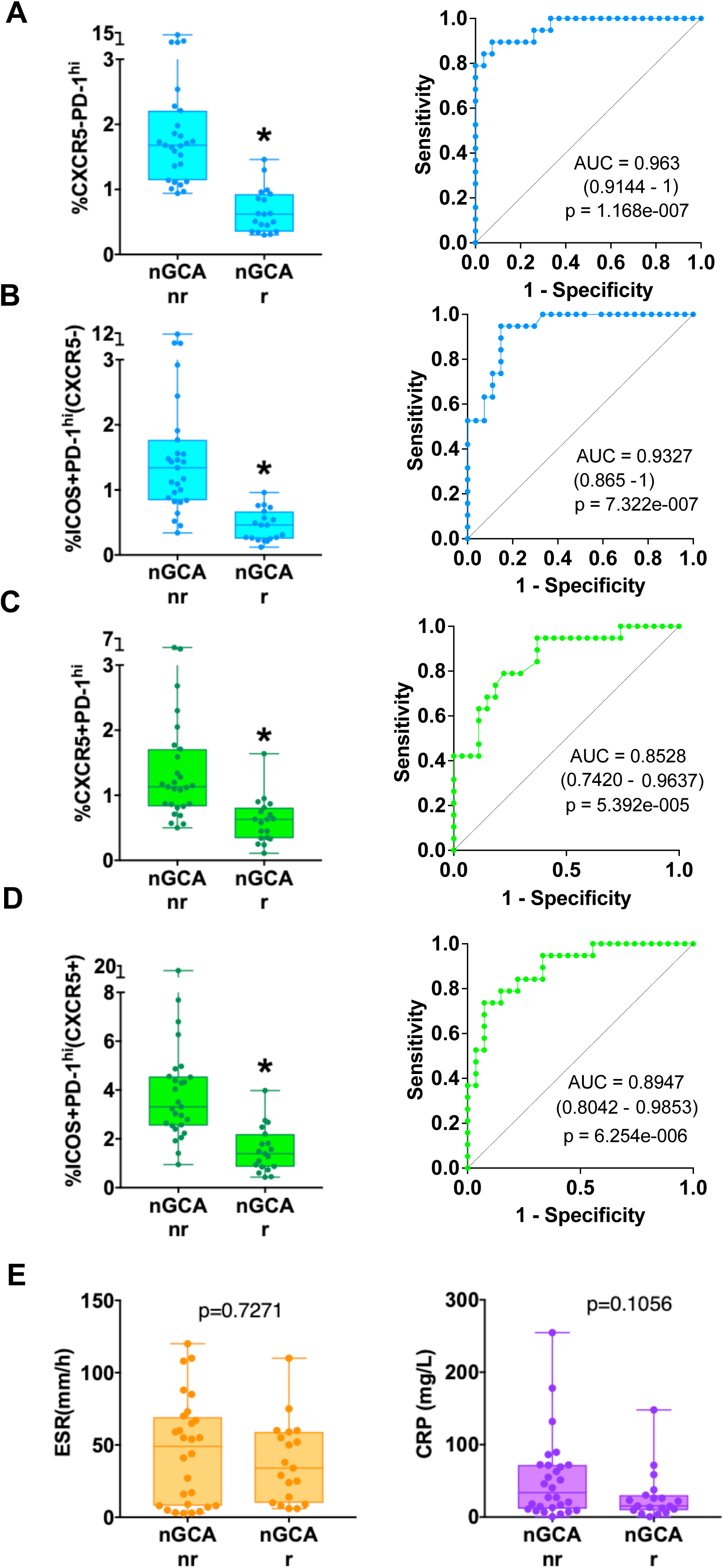
Relation of the occurrence of nGCA relapse in the first 12 months after diagnosis, with the baseline cTph and cTfh cell frequencies or with the baseline ESR and CRP values. **A-D**. Baseline cTph (A), ICOS+cTph (B), cTfh (C), and ICOS+cTfh (D) cell frequencies, in nGCA patients who were followed up clinically for 12 months and did not relapse (“nr”, n=27), vs nGCA patients who were followed for 12 months and relapsed (“r”, n=19). Bars in the left panels represent the median and interquartile range; whiskers represent the maximum and minimum values. *p<0.05, Mann-Whitney test. The right panels show the ROC curves representing the relation of the baseline cTph (A), ICOS+ cTph (B), cTfh (C) and ICOs+ cTfh (D) cell frequencies with the occurrence of relapse in the first year (ROC AUC and p values calculated with logistic regression analysis). **E**. Baseline ESR and CRP values in nGCA patients who did not relapse (nr) or who relapsed (r) in the first 12 months.

The baseline ESR and CRP values were not significantly different between patients who subsequently relapsed and those who did not (Fig.3E).

### Modifications of cTph and cTfh cell frequencies in nGCA along the disease course

Thirty-five nGCA patients donated blood for a second time 6 months after the initiation of treatment, together with the same 35 HC who had participated initially. These patients were receiving between 15 and 20 mg of prednisone and were clinically stable with no signs of activity. As compared with the baseline values, the frequencies of cTph, ICOS+cTph, cTfh and ICOS+cTfh cells had significantly decreased in nGCA patients but had not varied in HC (Fig 4 A-D). In three of these patients, blood was also drawn at the time of relapse; a rise in their cTph and cTfh cell frequencies was then observed, up to levels comparable to those observed at baseline (Fig. 4E).

**Figure 4.**
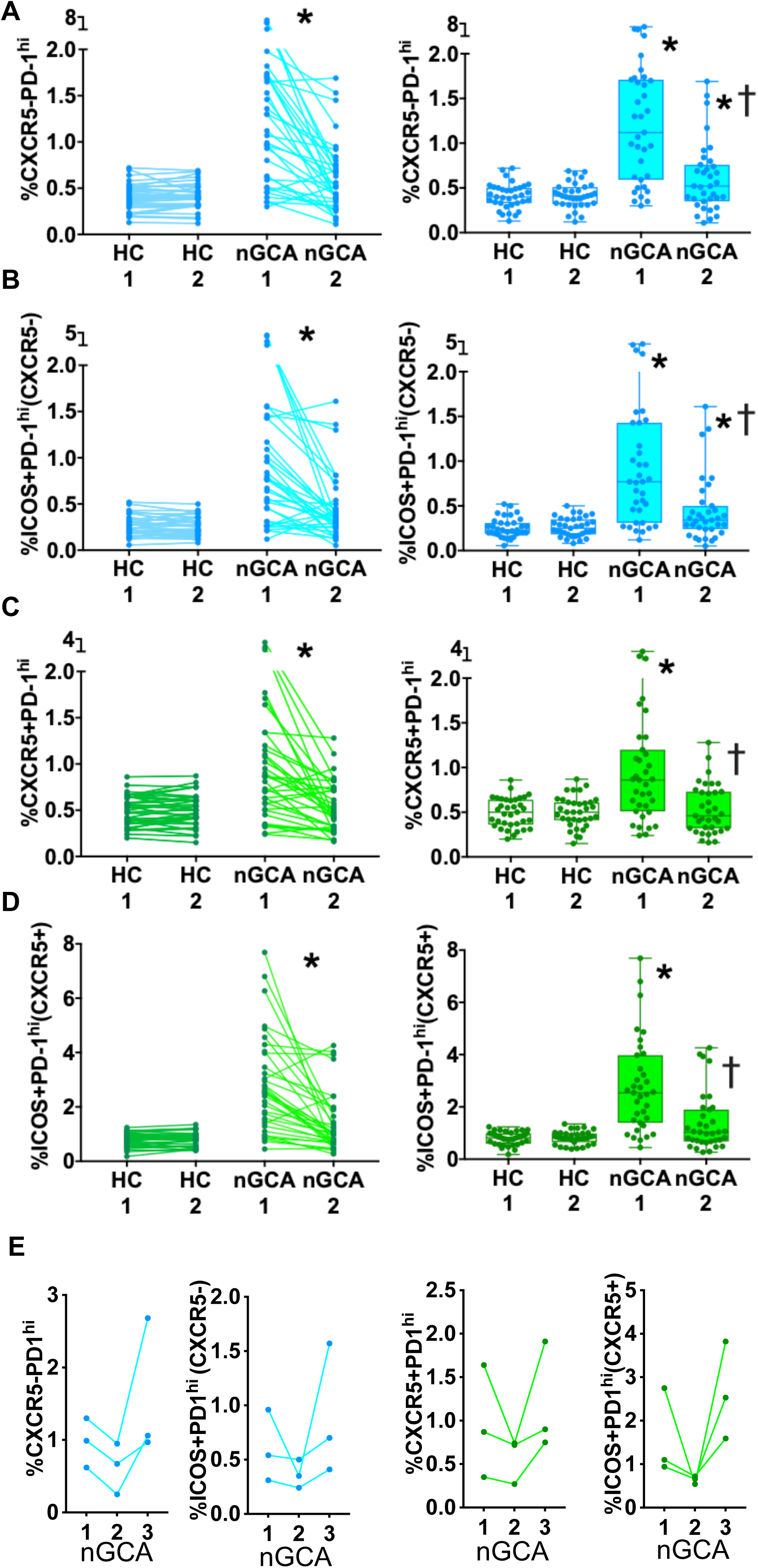
Modifications of cTph and cTfh cell frequencies in nGCA along the disease course. **A-D**. Values of cTph (A) ICOS+cTph (B), cTfh (C) and ICOS+cTfh (D) cell frequencies in nGCA (n=35) at baseline and 6 months after the initiation of glucocorticoid monotherapy, and in their untreated age and gender-matched HC (n=35). The left panels show paired longitudinal representations, p values calculated with Wilcoxon test. The right panels show unpaired representations, *p<0.05 vs HC, Kruskal-Wallis test with Bonferroni correction; † p<0.05 vs nGCA at baseline, Mann Whitney test. **E**. Values of cTph, ICOS+cTph, cTfh, and ICOS+cTfh cell frequencies in nGCA patients (n=3) at three different time points: baseline (“1”), clinically stable (“2”) and relapse (“3”).

## DISCUSSION

GCA, a granulomatous vasculitis typically affecting elderly patients, is a systemic condition associated with a substantial inflammatory response and a significant risk of severe clinical complications [1]. Despite intensive research its pathogenesis is not well understood and biomarkers for clinical relapse have not been clearly identified.

We are herein reporting for the first time that nGCA patients demonstrate increased baseline frequencies of two circulating CD4+ T cell populations expressing high PD-1 levels (CXCR5-/cTph and CXCR5+/cTfh); in addition, the activated ICOS+cTph and ICOS+cTfh cells were also increased. This points to their pathophysiological implication and interestingly, these frequencies associate with the clinical outcome. In fact, in patients who completed a 12 month follow-up period, separate analysis attending to the clinical outcome revealed that the baseline cTph and cTfh cell frequencies had been significantly higher in non-relapsing vs relapsing patients. This suggests that an insufficient ability to upregulate the checkpoint inhibitor PD-1 conditions a poorer prognosis, and is consistent with published results indicating the participation of the PD-1/PD-L1 system in GGA pathogenesis [2,4–6].

GCA has been considered as a T cell dependent condition [26,27], and an intense infiltrate of activated PD-1+ T cells is apparent at the inflamed vessel walls [2,3]. PDL-1+ monocytes and dendritic cells are also observed [2,3] although their level of PDL-1 expression seems to be insufficiently upregulated [2]. In addition, B lymphocytes and plasma cells are locally present [28,29] and an altered B cell profile has been described in the peripheral blood of GCA patients [30]. Furthermore, tertiary lymphoid structures have been observed in the GCA cranial arteries [28] and aorta [16,29].

CD4+CXCR5-PD-1^hi^ Tph cells were initially described in the inflamed synovium of seropositive RA patients [7] and contribute to the local inflammatory process. Subsequently, Tph cells have been related to additional autoimmune conditions [9] such as systemic lupus erithematosus (SLE) [31–33] Sjögreńs syndrome [34], ANCA-associated vasculitis (AAV) [35], type 1 diabetes mellitus [36], celiac disease [37] or ulcerative colitis (UC) [38]. However, to our knowledge they have not been previously reported in GCA. Because Tph cells are typically located at inflammatory foci [7], they may account for the PD-1+ cells observed at the GCA arterial wall infiltrates [2,3]. Our current study is limited by the lack of GCA biopsy analysis. This limitation is partially compensated since circulating Tph cells (cTph) can be detected in the peripheral blood [7–9, 31–37] and previous reports by other investigators have elegantly demonstrated the local presence of PD-1+T cells [2,3]. In patients with seropositive RA, SLE, UC or AAV, the frequency of cTph and ICOS+cTph varies in parallel with changes of disease activity [7,8,31–33,35,38]. Accordingly, we observed that their baseline levels in our nGCA patients decreased after the initiation of treatment, in association with clinical improvement. Furthermore, these frequencies increased again at the time of relapse.

Contrasting with the characteristic presence of Tph at inflammatory foci, Tfh cells are typically located in the germinal centers of secondary lymphoid organs [10,11], where they promote B cell differentiation and function [10,11]. Tfh cells are not found in the diffuse lymphoid infiltrates at inflamed sites [7,39] but have been observed in ectopic lymphoid structures [28,29,16], to which Tfh may be recruited by CXCL13-secreting Tph cells [9]. In addition, circulating Tfh-like cells are observed in the peripheral blood and have been termed “circulating Tfh” or “cTfh” [10,11]; this facilitates the study of this population in human subjects, given the cumbersome access to secondary lymphoid organs. Circulating Tfh cells are Bcl-6 negative, express CXCR5, PD-1 and/or ICOS [10,11], provide help to B cells [10,11] and their frequencies are elevated in patients with several autoimmune conditions [8–15].

Previous studies of cTfh cells in GCA are scant and use different cytometry gating strategies to define this population [16,17]. Matsumoto et al. [17] and Desbois et al. [16] defined cTfh in GGA and Takayasu arteritis patients as CD4+CXCR5+ T cells and did not study their PD-1 or ICOS expression. In fact, the criteria used for defining cTfh cells are discordant among published studies [10,12,16,17,40]. Certainly, the population of circulating CD4+CXCR5+PD-1^hi^ and/or ICOS+ T cells is small [10] and can only be adequately detected when examining a large number of CD4+ T cells by cytometry. Some authors describe cTfh as CD4+CXCR5+ [16,17] or CD4+CXCR5+PD-1+ [13,15]. Yet, other investigators consider that PD-1^lo^CXCR5+ CD4+ T cells can be central memory T cells, also observed in the circulation of healthy individuals [12, 40] whereas PD-1^hi^CXCR5+ CD4+ T cells represent an active Tfh cell differentiation in lymphoid organs and can be used to monitor the activation status of the Tfh program [41].

Matsumoto et al. [17] reported that circulating CD4+CXCR5+ T cells are elevated at baseline in Takayasu but not in GCA; their frequencies decreased in parallel with clinical improvement in both conditions and rised again in patients who relapsed [17]. Likewise, Desbois et al. observed augmented CD4+CXCR5+ T cells in the peripheral blood of Takayasu but not GCA [16] and also reported an increased circulating Tfh17 (CD4+CXCR5+CCR6+) cell signature in patients with Takayasu arteritis that was not present in GCA [16]. Consistent with these observations we did not find differences in the frequency of circulating CD4+CXCR5+ T cells between GCA patients and HC. To our knowledge there are no previous reports indicating the frequency of CD4+CXCR5+/PD-1+ or PD-1^hi^ cTfh cells in GCA. Hid Cadena et al. [3] observed a reduced frequency of circulating CD4+PD-1+ T cells in patients with GCA but did not discriminate between CXCR5+ or CXCR5-. Theirs was a cross-sectional study of patients with diverse clinical activity levels under various treatment regimes [3] and contrasts with the elevated cTfh and cTph cell frequencies we are reporting in untreated nGCA; this discordance may be explained by differences in study design and clinical features.

Contrasting with the transient elevation of cTfh cells we are herein reporting in nGCA, we previously published that patients with seropositive RA demonstrate a constitutive elevation of cTfh cells, which is independent of the presence or absence of disease activity [8,14]. This suggests that the pathogenic implication of Tfh cells in nGCA is different from RA and may be related to the lack of detection of autoantibodies in this condition. In fact, altered Tfh cell numbers have been observed in other autoimmune disorders that do not seem to be mediated by autoantibodies [11], implying that the role of Tfh cells in autoimmunity may not be limited to supporting autoantibody generation [11].

Interestingly we did not find significant differences in cTph or cTfh cell frequencies among ultrasound-defined subgroups. Indeed, a distinction was initially recognized separating cranial and extracranial phenotypes [25]; however, the task force subsequently acknowledged that GCA constitutes a single entity with a range of overlapping phenotypes [25,42]. In fact, around 35% of the patients presenting with classical “cranial” clinical manifestations demonstrate large vessel involvement in imaging studies [43]. Also, patients classified at disease onset as large vessel GGA may develop clinical and US cranial involvement at follow-up [44,45], and viceversa. In addition, we observed that the frequencies of cTph, ICOS+cTph, cTfh or ICOS+cTfh were not significantly different between patients with or without PMR.

In the present study, our patients received treatment according to the current EULAR recommendations [18]. These are based on the observation that a significant proportion of patients with GCA receiving GC monotherapy do not relapse and are able to reduce the GC dose down to ≤5 mg/day after 1 year [18,46–48]; hence, the prompt initiation of DMARDs in steroid-responsive subjects would unnecessarily promote safety hazards [18] and increase costs [18,49]. Our results on the predictive value of cTph and cTfh cells may help identify patients with an increased risk of relapse who would benefit from an early treatment intensification and/or addition of DMARDs. It also suggests that patients with low PD-1 expression could demonstrate a good response to PD-1 agonistic therapies. We did not observe an association of the baseline ESR or CRP values with the clinical outcome, although these parameters demonstrated a weak correlation with the baseline cTph and/or cTph cell frequencies. Previous studies have yielded conflicting results regarding the predictive value of ESR or CRP in GCA [50]; discrepancies may be related to different study designs, treatment regimes and/or clinical features.

In summary, the transiently elevated frequencies of cTph and cTfh cells in nGCA suggests their implication in the pathogenesis of this condition and may constitute a useful biomarker for personalized therapeutic strategies, thereby contributing to minimizing the disease associated burden.

## Key messages

### What is already known on this topic

- The PD-1/PD-L1 coinhibitory pathway is essential for restraining autoimmunity
- Therapeutic inhibition of this pathway has been associated with new onset of giant cell arteritis (GCA)
- Activated PD-1+ T cells have been observed in GCA arterial wall infiltrates
- The PD-1^hi^ Tph and Tfh cell frequencies are elevated in the peripheral blood of patients with several autoimmune conditions but have not been investigated in GCA
- In clinical practice we lack risk markers for GCA relapse.

### What this study adds

- This is the first description indicating that patients with treatment-naïve, newly diagnosed GCA demonstrate increased baseline frequencies of circulating PD-1^hi^Tph (cTph) and Tfh (cTfh) cells, and that
- The baseline cTph and cTfh cell frequencies associate with the occurrence of relapse in the first 12 months after diagnosis in patients treated with glucocorticoid monotherapy

### How this study might affect research, practice or policy

- The cTph and cTfh cell frequencies may help identify nGCA patients with an increased risk of relapse who would benefit from an early treatment intensification and/or addition of DMARDs.
- These results also suggests that patients with low PD-1 expression could demonstrate an optimal response to PD-1 agonistic therapies.
- Therefore, the cTph and cTfh cell frequencies are potential tools for personalized medicine strategies in nGCA.

---

**Figure S1.**
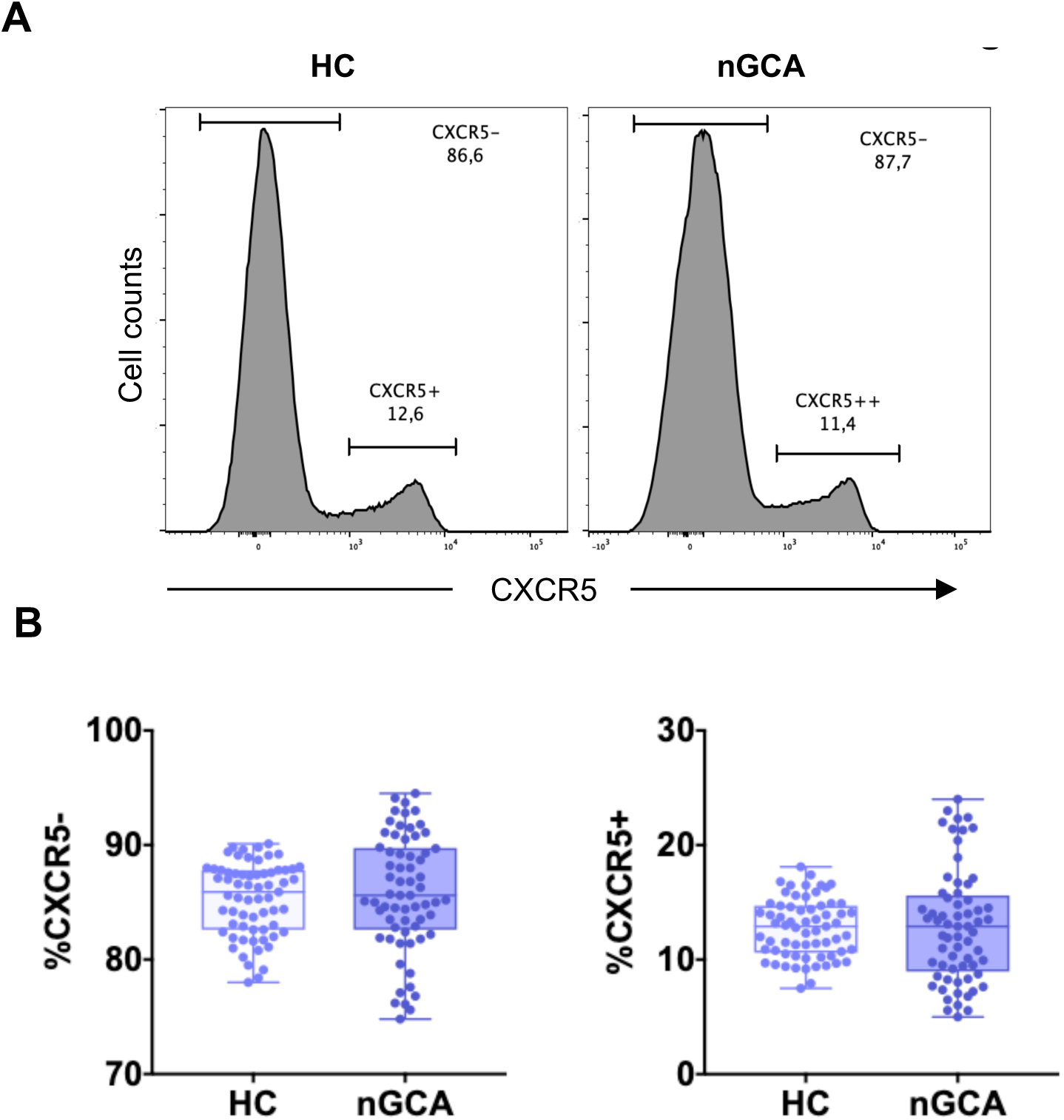
Baseline frequency of CXCR5- and CXCR5+ cells among circulating CD4 + T lymphocytes. **A**. Representative flow cytometry histograms of CXCR5 expression in cells gated for CD3 and CD4 expression. **B**. Baseline frequency of CXCR5- (left) and CXCR5+ (right) cells among CD4 T lymphocytes in nGCA patients (n=65) and HC (n=65). Box and whiskers plots represent the median, interquartile range, maximum and minimum values.

